# Robustness of a multivariate composite score when evaluating distress of animal models for gastrointestinal diseases

**DOI:** 10.1101/2022.11.14.516441

**Authors:** Steven R. Talbot, Simone Kumstel, Benjamin Schulz, Guanglin Tang, Ahmed Abdelrahman, Nico Seume, Edgar Heinz Uwe Wendt, Johanna Eichberg, Christine Häger, Andre Bleich, Brigitte Vollmar, Dietmar Zechner

## Abstract

The fundament of an evidence-based severity assessment in laboratory animal science is reliable distress parameters. Many readouts are used to evaluate and determine animal distress and the severity of experimental procedures. Therefore, we analyzed four distinct parameters like the body weight, burrowing behavior, nesting, and distress score in the four gastrointestinal animal models (pancreatic ductal adenocarcinoma (PDA), pancreatitis, CCl_4_ intoxication, and bile duct ligation (BDL)). Further, we determined the parameters‘ robustness in various experimental subgroups due to slight variations like drug treatment or telemeter implantations. We used non-parametric bootstrapping to get robust estimates and 95 % confidence intervals for the experimental groups. It was found that the performance of the readout parameters is model-dependent and that the distress score is prone to experimental variation. On the other hand, we also found that burrowing and nesting can be more robust than, e.g., the body weight when evaluating PDA. However, the body weight still was highly robust in BDL, pancreatitis, and CCl_4_ intoxication. To address the complex nature of the multi-dimensional severity space, we used the Relative Severity Assessment (RELSA) procedure to combine multiple distress parameters into a score and mapped the subgroups and models against a defined reference set obtained by telemeter implantation. This approach allowed us to compare the severity of individual animals in the experimental subgroups using the maximum achieved severity (RELSA_max_). With this, the following order of severity was found for the animal models: CCl_4_ < PDA ≈ Pancreatitis < BDL. Furthermore, the robustness of the RELSA procedure and outcome was externally validated with a reference set from another laboratory also obtained from telemeter implantation. Since the RELSA procedure reflects the multi-dimensional severity information and is highly robust in estimating the quantitative severity *within* and *between* models, it can be deemed a valuable tool for laboratory animal severity assessment.

## Introduction

Laboratory animals have made significant contributions to biomedical research [1–4]. However, public and political concerns regarding experiments on animals are steadily increasing [5–8]. Several legislative advancements have been made to alleviate these concerns and to expand animal welfare taking scientific requirements, ethics, and morals into account. These advancements led to the broad adoption of the 3R principles and the passing of the European Union Directive 2010/63/EU [9,10]. Articles 38, 39, 54, and the Annex VIII of Directive 2010/63/EU demand prospective and retrospective assessment of the severity of experimental procedures, classified into four categories (mild, moderate, severe, and non-recovery) [11].

Similarly, the USA’s Institutional Animal Care and Use Committees enforce the local Animal Welfare Act and Animal Welfare Regulations [12]. China, estimated to be the top user of animals for experimental research, has also implemented guidelines for ethical review of laboratory animal welfare in late 2018 [13]. Consequently, appropriate methods to assess severity and distress in animal research are of utmost importance. Although significant improvement has been made in regards to the refinement of procedures, such as appropriate analgesia [14] and humane endpoints [15], the lack of validated, evidence-based methodology hinders the advancement of animal welfare as well as the corresponding science [16,17].

Several readout parameters for animal distress evaluation have been found in recent years. In addition, researchers often use non-invasive methods to assess physical and physiological parameters, appearance, and behavior. For example, body weight is widely used to determine animal well-being and to refine humane endpoints in experimental procedures [18]. Score sheets are also routinely used in judging clinical signs to assess the level of distress in animals. Such scores often provide the basis for subsequent actions if predefined humane endpoints are reached [19,20]. Additionally, evaluating innate behavior like burrowing and nesting activity has become a cornerstone of assessing animal well-being, as they are reduced in severe distress [21–23].

Although it is well accepted that multiple rather than single readout parameters should be used to describe and compare animal distress [24–26], single parameters are often not combined to form a single score [27–30]. However, a combined analysis of multiple readout parameters is vital to explore the multi-dimensional dependencies of the variables. Depending on the nature of the analysis, multivariate or multiple-variable approaches like regressions are used [31]. Publications have shown that multivariate methods such as Principal Component Analysis (PCA) (Ernst et al. 2020; Häger et al. 2018) and *k*-means clustering [32,33] can be used to evaluate the performance and the importance of variables in animal models, i.e., as indicators of animal distress. However, methods like PCA are prone to collinearity – which often occurs in the measured variables, and, therefore, require careful analysis. More sophisticated strategies involve, e.g., Machine Learning [34] and adaptive modeling [35] to assess and select individual variables or combinations. The omnipresent high variance in animal experiments is not only a problem regarding reproducibility but also hampers the usefulness of statistical methods. Non-parametric methods like bootstrapping [36,37] can help to obtain more reliable parameter estimators and the corresponding confidence intervals [38]. This method was deemed superior to classical inferential statistics, especially in the clinical context [39,40].

While all readout parameters mentioned above proved helpful in the past, little is known about their robustness in different experimental settings. Robustness, in a broader scientific sense, means that conclusions remain stable even when experimental conditions are varied [41,42]. Furthermore, robustness has been defined as one of three critical aspects of the usefulness of an animal experiment [43,44]. Thus, it is essential for translational research and ethical considerations when judging specific animal models. For example, suppose an animal model’s distress can be measured robustly. In that case, it makes sense to argue in favor or against that specific model to improve animal well-being by refining interventions and procedures.

Consequently, one purpose of this study was to evaluate the robustness of animal distress parameters (burrowing activity, nesting behavior, body weight, distress score) when varying experimental conditions. In addition, it was the goal to combine these readout parameters into a single metric, called RELSA_max_ [45] and to evaluate whether this score can be used to differentiate the distress levels of four animal models for gastrointestinal diseases (pancreatic cancer, chronic pancreatitis, liver fibrosis, and cholestasis). Finally, this study aimed to check the robustness of the RELSA_max_ score by employing different sets of reference data from independent laboratories.

## Material and Methods

### Animal models

#### Animals

This study did not use new animals but re-evaluated data generated in previous projects with the novel focus of combining multiple distress parameters into a score to compare the severity of various animal models. All animal experiments were approved by the local authority (Landesamt für Landwirtschaft, Lebensmittelsicherheit und Fischerei Mecklenburg-Vorpommern (license 1-019/15, 1-062/16, 1–002/17) or the Lower Saxony State Office for Consumer Protection and Food Safety (LAVES, license 15/1905). Mice in laboratory A were housed at the central animal facility of the University Medical Center Rostock in different type III cages (Zoonlab GmbH, Castrop-Rauxel, Germany) at 12 h light/dark cycle (light period: 7:00–19:00), a temperature of 21 ± 2 °C, and relative humidity of 60 ± 20 % with food (10 mm pellets, ssniff-Spezialdiäten GmbH, Soest, Germany) and tap water ad libitum. Enrichment was provided in the form of a paper roll (75 × 38 mm, H 0528–151, ssniff-Spezialdiäten GmbH), nesting material (shredded tissue paper, Verbandmittel GmbH, Frankenberg, Deutschland), and a wooden stick (40 × 16 × 10 mm, Abedd, Vienna, Austria). The health of the animal stock was routinely checked according to FELASA guidelines (Helicobacter sp., Rodentibacter pneumotropicus, and murine Norovirus were detected in a few mice within the last two years; these animals were not used for any experiments).

Mice for the reference data set B were pair-housed at the Central Animal Facility of the MHH in macrolon type-II cages (360 cm^2^; Tecniplast, Italy), which were changed once per week. Cages were bedded with autoclaved softwood shavings (poplar wood; AB 368P, AsBe-wood GmbH, Buxtehude, Germany), paper nesting material (AsBe-wood GmbH, Buxtehude, Germany), and two cotton nesting pads (AsBe-wood GmbH, Buxtehude, Germany). Room conditions were standardized (22 +/- 1°C; humidity: 50 %-60 %; 14:10 h light/dark cycle). Mice were fed standard rodent food (Altromin 1324, Altromin, Lage, Germany) *ad libitum*, and autoclaved (135°C/60 minutes) distilled water was provided *ad libitum*. All mice were randomly allocated to the experimental groups and habituated to the experimental environment before the surgical procedure. The mice were free of the viral, bacterial, and parasitic pathogens listed in the recommendations of the Federation of European Laboratory Animal Science Association.

ETA-F-10 transmitters (Data Sciences International, Minnesota, USA) in laboratory A were placed in the abdominal cavity of male C57BL/6J mice after anesthetizing them with 1–2 vol % isoflurane (n= 10). Analgesia was provided by one s.c. injection of 5 mg/kg carprofen (Rimadyl, Pfizer GmbH, Berlin, Germany) before surgical intervention and 1250 mg/L metamizole (Ratiopharm, Ulm, Germany) in the drinking water until the end of the experiment. This experiment’s methodological details and data were published previously [24,46].

In laboratory B, transmitters (ETA-F10 or HD-X11; DSI, St Paul, MN, USA) were aseptically implanted into the intraperitoneal cavity of 9-10 weeks old female mice C57BL/6J (n=13) with electrodes placed subcutaneously for a bipolar lead II configuration under general isoflurane anesthesia. General anesthesia was induced in an induction chamber (15 × 10 × 10 cm) with 5 vol % isoflurane (Isofluran CP®, CP Pharma, Burgdorf, Germany) and an oxygen flow (100 % oxygen) of 6 l/min. After confirmation of the absence of the righting reflex and removal from the chamber, anesthesia was maintained via an inhalation mask with 1.5-2.5 vol % isoflurane and an oxygen flow of 1 l/min. The corneal reflex was combined with the eyelid-closing reflex and the toe pinch reflex to determine the depth of anesthesia. Personnel involved have been trained and were experienced in performing these assays carefully and very softly to omit any damage. In addition, the eyes were moistened with eye ointment to protect them from drying (Bepanthen®, Bayer AG, Leverkusen, Germany). After total anesthesia, the surgical area was shaved, and the mice were placed in the surgical field in dorsal recumbency with the head towards the surgeon. During the entire duration of the anesthesia, the mice were placed on a heating pad at 37.0 ± 1.0°C to prevent hypothermia. EMLA creme (1g of cream = 25 mg/g Lidocaine + 25 mg/g Prilocaine; Aspen Germany GmbH, Munich, Germany) was used for local anesthesia at the incision sites. For analgesia, animals received either preoperative 200 mg/kg metamizole (Novaminsulfon 500 mg Lichtenstein, Zentiva Pharma GmbH, Frankfurt am Main, Germany) subcutaneously (s.c.) and postoperative 200 mg/kg metamizole orally via the drinking water until day 3 or preoperative 5 mg/kg carprofen (Rimadyl, Zoetis Deutschland GmbH, Berlin, Germany) s.c. and postoperative 2.5 mg/kg s.c. every 12 h until day 3. The methodological details and the data of this experiment were published previously [45].

Pancreatic cancer was established by injecting 2.5 × 10^5^ 6606 PDA cells slowly into the pancreas of anesthetized male C57BL/6J mice. For all mice, analgesia was provided by one s.c. injection of 5 mg/kg carprofen (Rimadyl, Pfizer GmbH) before cell injection and 1250 mg/L metamizole (Ratiopharm) in the drinking water until the end of the experiment. Mice (n=7) of one subgroup had an ETA-F-10 transmitter implanted 14 days before cell injection (Kumstel et al. 2020b). All other mice had no transmitter [47]. Starting on day 4 after cell injection, mice without transmitters were treated with combinatorial chemotherapies or the appropriate vehicles as controls (vehicle CHC/Met: n = 7, vehicle Gal/Met: n = 5). Either α-cyano-4-hydroxycinnamate (CHC, daily i.p. injection of 15 mg/kg, Tocris Bioscience, Bristol, UK) plus metformin (Met, daily i.p. injection of 125 mg/kg, Merck, Darmstadt, Germany) or galloflavin (Gal, i.p. injection of 20 mg/kg three times a week, Tocris Bioscience) plus metformin (Met, daily i.p. injection of 125 mg/kg, Merck) were applied as chemotherapies until day 37 after cell injection (CHC/Met: n= 7, Gal/Met: n = 7). This experiment’s methodological details and data were published previously [24,47].

Male C57Bl/6J mice were treated with cerulein (Bachem H-3220.0005, Bubendorf, Switzerland) to induce chronic pancreatitis. Cerulein was dissolved in 0.9 % sodium chloride and administered by consecutive intraperitoneal (i.p.) injections (50 μg/kg, three hourly injections/day on three days/week for four weeks. MicroRNA-21 inhibitor (miRCURY LNA™ microRNA-21a-5p inhibitor; sequence: TCAGTCTGATAAGCT) and its corresponding microRNA-21 control (miRCURY LNA™ microRNA-21a-5p control; sequence: TCAGTATTAGCAGCT) were purchased from Qiagen (Hilden, Germany), resuspended in PBS and injected at a dose of 10 mg/kg (s.c.) on day 0 and day 14 after first cerulein injection (inhibitor: n = 8, vehicle: n = 8). This experiment’s methodological details and data were published elsewhere [24,48].

For inducing liver damage, carbon tetrachloride (Merck Millipore, Eschborn, Germany) was diluted fourfold with corn oil (Sigma-Aldrich, code C8267), and 1 μl per g body weight of this solution (dosage: 0.25 ml/kg body weight) was injected (i.p.) into male BALB/cANCrl mice twice per week over six weeks. Analgesia was provided by 1250 mg/L metamizole (Ratiopharm) in the drinking water until the end of the experiment. 20 mg/kg MCC950 (Sigma Aldrich, St. Louis, USA) or aqua dest. ad inj. (vehicle control) was injected (i.p.) into mice (for nesting activity MCC950: n = 6, vehicle: n = 6; for burrowing MCC950: n = 7, vehicle: n = 3; for body weight and distress score: MCC950: n = 13, vehicle: n = 9) daily from day 28 to day 41 after the first carbon tetrachloride injection. This experiment’s methodological details and data were published elsewhere [24,49].

A laparotomy was performed on male BALB/cANCrl mice under anesthesia (1.2–2.5 vol % isoflurane) to induce cholestasis by bile duct ligation (BDL). The bile duct was ligated by three surgical knots and transected between the two distal ligations. To relieve pain, 5 mg/kg carprofen (Pfizer GmbH, Berlin, Germany) was injected (s.c.) before the operation, and 1250 mg/L metamizole (Ratiopharm) was provided in the drinking water until the end of the experiment. 20 mg/kg MCC950 or aqua dest. Ad inj. (vehicle control) was i.p. injected into mice (for nesting activity MCC950: n = 7, vehicle: n = 7; for burrowing and distress score MCC950: n = 7, vehicle: n = 9; for body weight: MCC950: n = 14, vehicle: n = 16) daily from day 1 before BDL to day 13 after BDL. This experiment’s methodological details and data were published elsewhere [24,49].

### Evaluation of distress

The body weight, burrowing activity, nesting, and distress score were evaluated to assess distress. In laboratory A, the burrowing activity was analyzed by filling a tube (length: 15 cm, diameter: 6.5 cm) with 200 g of food pellets, which was then placed into the mouse cage 2–3 h before the dark phase. The remaining pellets in the burrowing tube were weighed after 2 hours (for C57Bl/6J mice) or 17 ± 2 h (for BALB/cANCrl mice), and the weight of the burrowed pellets was calculated. The percentage of burrowing activity and body weight was calculated by using the weight of burrowed pellets or body weight before any intervention as a reference.

The nest-building behavior in laboratory A was analyzed by placing a cotton nestlet (5 cm square of pressed cotton batting, Zoonlab GmbH, Castrop-Rauxel, Germany) in the cage 30 to 60 minutes before the dark phase. The nests were scored at the end of the dark phase ± 2 hours using a scoring system developed by Deacon [50]. However, a 6^th^ score point was added to this scoring system. A score of 6 defined a perfect nest: The nest looked like a crater, and more than 90 % of the circumference of the nest wall was higher than the body height of the coiled-up mouse. Please note that the nesting activity of BALB/cANCrl mice was measured 1 day after evaluating the burrowing activity to avoid offering the animals two actions at the same time.

In addition, the well-being of mice was assessed at laboratory A by evaluating multiple parameters with the help of a scoresheet. This scoresheet was based on other score sheets [19,20] and was previously published by our group [51]. The mice were, therefore, observed in their home cage for a few minutes, and the distress score was assessed when one or more defined criteria (e.g., spontaneous behavior, flight behavior, or general body conditions) were diagnosed.

Data obtained from laboratory B served as the reference set. Here, animal distress was assesses by analysis of burrowing behavior. Here, baseline measurements were taken on days two and one before surgery. A 250 mL plastic bottle with a length of 15 cm, a diameter of 5.5 cm, and a port diameter of 4 cm was used as a burrowing apparatus. It was filled with 140 g +/- 1.5 g of the standard diet pellets of the mice (Altromin1324, Lage, Germany). On day 1, 2, 3, 5, and 7 after surgery, mice were singly placed in a type-II macrolon cage with autoclaved hardwood shavings overnight. The burrowing bottles were placed in the left corner. Half of the used nesting material from the home cage was provided as a shelter in the right corner. The tests started three hours before the dark phase. The amount of burrowed pellets was assessed after two hours and after 12 hours. Body weight were evaluated two days before (baseline) and daily after transmitter implantation (for one week). This experiment’s methodological details and data were published elsewhere [45].

### Statistics

All statistical analyses were performed using the R software (v4.0.3) [52]. The continuous variables (body weight and burrowing activity) were standardized to 100 % at baseline levels (day=-1). Variables representing animal distress (body weight, burrowing activity, nesting score, and the distress score; Figure 1-4) were bootstrapped 10000-fold to obtain assumption-free estimates on the median (ŷ) and the 95 % confidence intervals (rcompanion [53]). In addition, the distribution of the estimates was inspected visually and tested against the hypothesis of normal distribution using Shapiro-Wilk’s test. In case of evidence for non-normally distributed data, the experimental subgroups were compared using the Kruskal-Wallis test. Pairwise comparisons were calculated with the Wilcoxon-Mann-Whitney test. Holm’s correction adjusted the resulting p-values for multiple comparisons, i.e., to control the family-wise error rate. Time-dependent (intra-treatment comparisons) of non–parametric data were analyzed with the Friedman Rank Sum test. Subsequent baseline-level comparisons were calculated with Dunn’s post hoc test and Holm’s correction. In the case of normally distributed data, a one-way ANOVA or repeated-measures ANOVA with sphericity corrections [54] for time-dependent data with “day” as the within-subjects variable was performed. Between-treatment analyses and comparisons to baseline levels (with the control group as day -1) were performed with Dunnett’s test and the Holm correction. Results were considered statistically significant at the following levels: * p≤0.05, **p≤0.01, ***p≤0.001, ****p≤0.0001; multiplicity-adjusted p-values are shown as p_adj_.

**Fig. 1:**
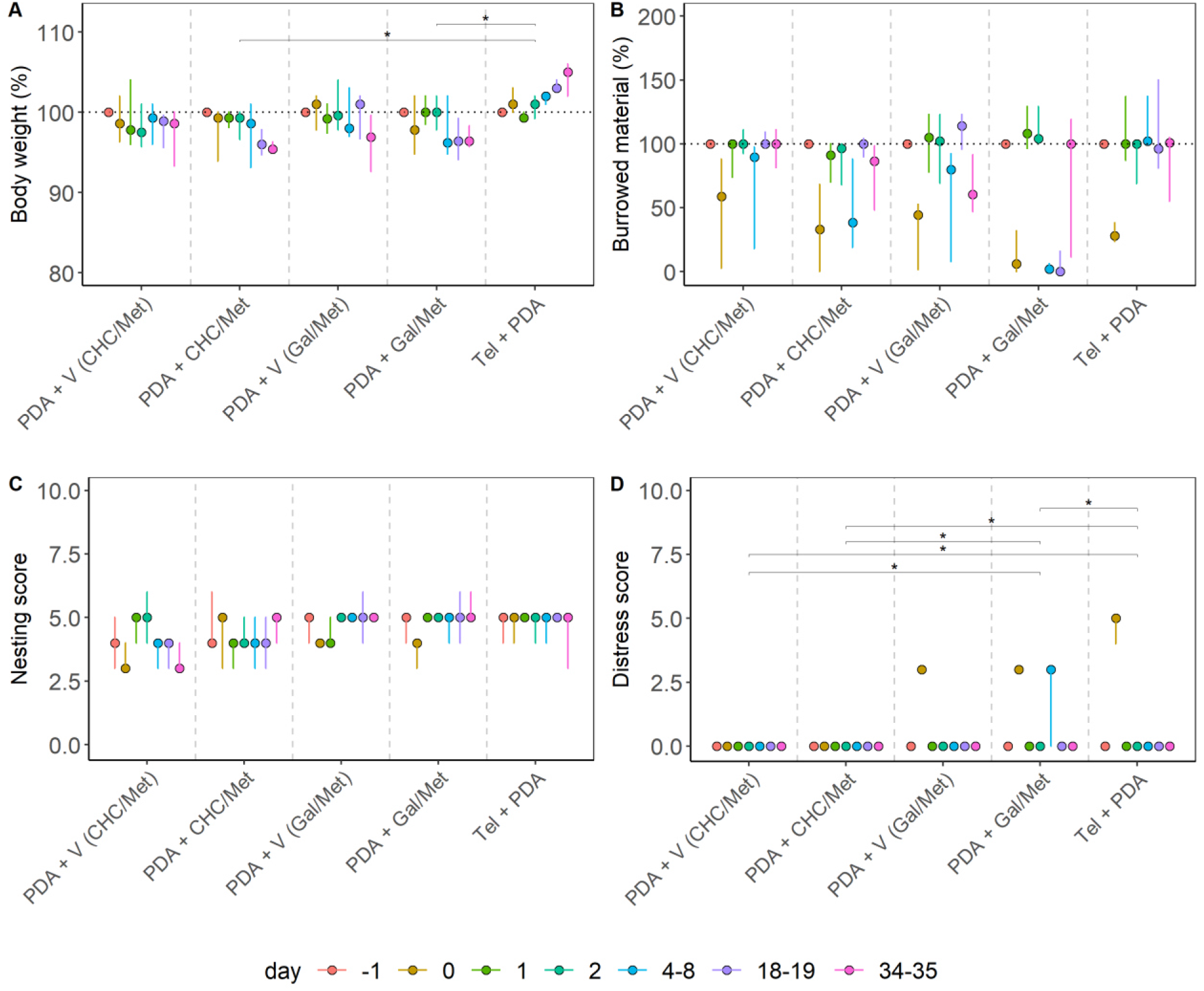
Distress evaluation of an orthotopic pancreatic cancer model. Pancreatic cancer (PDA) was treated with α-cyano-4-hydroxycinnamate plus metformin (CHC/Met), Galloflavin plus Metformin (Gal/Met), or the respective vehicle solutions (V). Telemetric transmitters were implanted into mice in a separate experiment, and 6606PDA cells were injected into the pancreas (Tel + PDA). The percentage of body weight (A), the percentage of burrowing activity (B), nesting (C), and the distress score (D) were evaluated on the indicated days. Differences between the groups in (A) were assessed with the Kruskal-Wallis test. Significant differences in body weight between the groups were detected (χ^2^ = 44.871, df = 4, p<0.0001), and the following post hoc comparisons revealed differences between the groups PDA + CHC/Met and Tel + PDA (*p_adj_ = 0.035) as well as between PDA + Gal/Met and Tel + PDA (*p_adj_ = 0.035). Group differences were also present in the distress score (χ^2^ = 12.67, df = 4, p=0.013), between PDA + V (CHC/Met) and PDA and Gal/Met (*p_adj_=0.015), PDA + V (CHC/Met) and Tel + PDA (*p_adj_=0.031), PDA + CHC/Met and PDA + Gal/Met(*p_adj_=0.015), PDA + CHC/Met and Tel + PDA (*p_adj_=0.031) as well as between PDA + Gal/Met and Tel + PDA (*p_adj_=0.031). The graphs depict the bootstrapped median estimator on each experimental day and the 95 % confidence intervals.

**Fig. 2:**
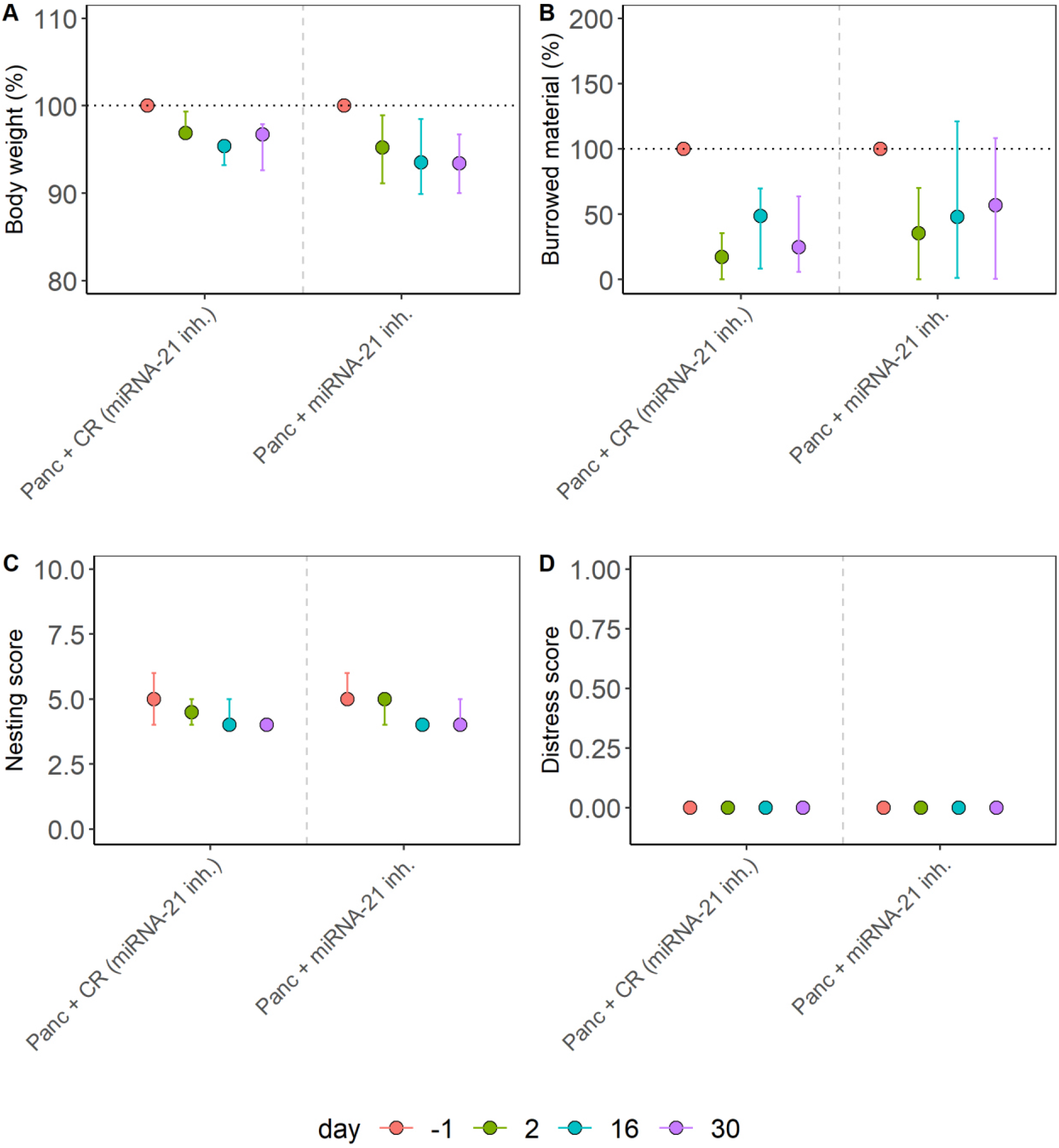
Distress evaluation of a chronic pancreatitis model. Chronic pancreatitis (Panc) was treated with a microRNA-21 inhibitor (miRNA-21 inh.) or the identical vehicle solution plus a respective control oligonucleotide (CR). The percentage of body weight (A), the percentage of burrowing activity (B), the nesting (C), and the distress score (D) were assessed on the indicated days. No significant differences between the two groups were determined using the Kruskal-Wallis test in A (χ^2^= 0.82, df = 1, p = 0.36), B (χ^2^=0.97, df = 1, p = 0.32), C (χ^2^= 0.45, df = 1, p = 0.5) or D (χ^2^= 0.44, df = 1, p = 0.50). The graphs depict the bootstrapped median estimator on each experimental day and the 95% confidence intervals.

**Fig. 3:**
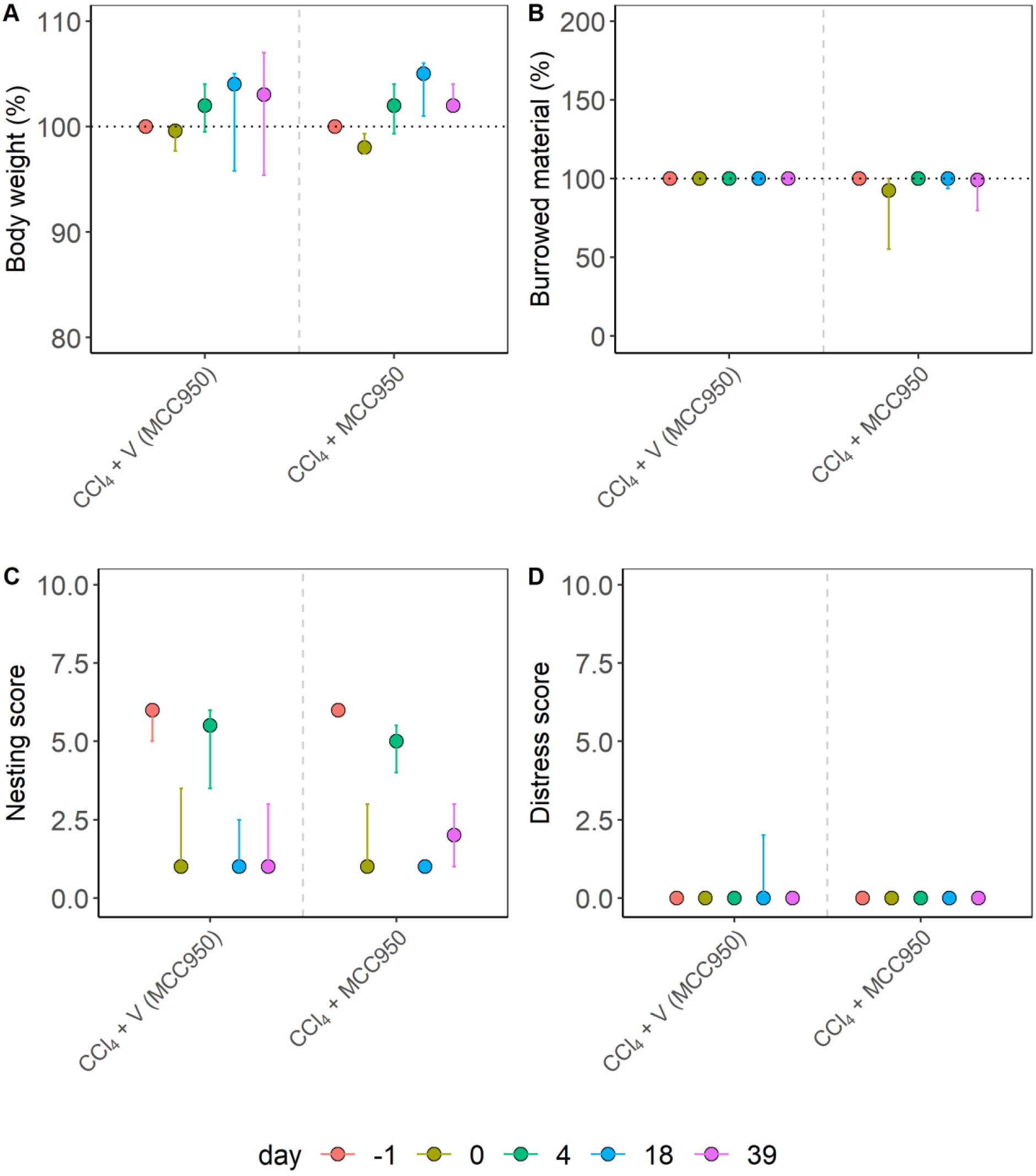
Distress evaluation of liver damage in a fibrosis model. Liver damage was induced by repetitive carbon tetrachloride application (CCl_4_), and mice were treated with an NLRP3 inflammasome inhibitor (MCC950) or the appropriate vehicle solution (V). The percentage of body weight (A), the percentage of burrowing activity (B), the nesting (C), and the distress score (D) were evaluated on the indicated days. No significant differences between the groups were found using the Kruskal-Wallis test in A (χ^2^= 0.48, df = 1, p = 0.49), B (χ^2^= 1.33, df = 1, p = 0.25), C (χ^2^= 0.047, df = 1, p = 0.83) or D (χ^2^= 2.33, df = 1, p = 0.13). The graphs depict the bootstrapped median estimator on each experimental day and the 95 % confidence intervals.

**Fig. 4:**
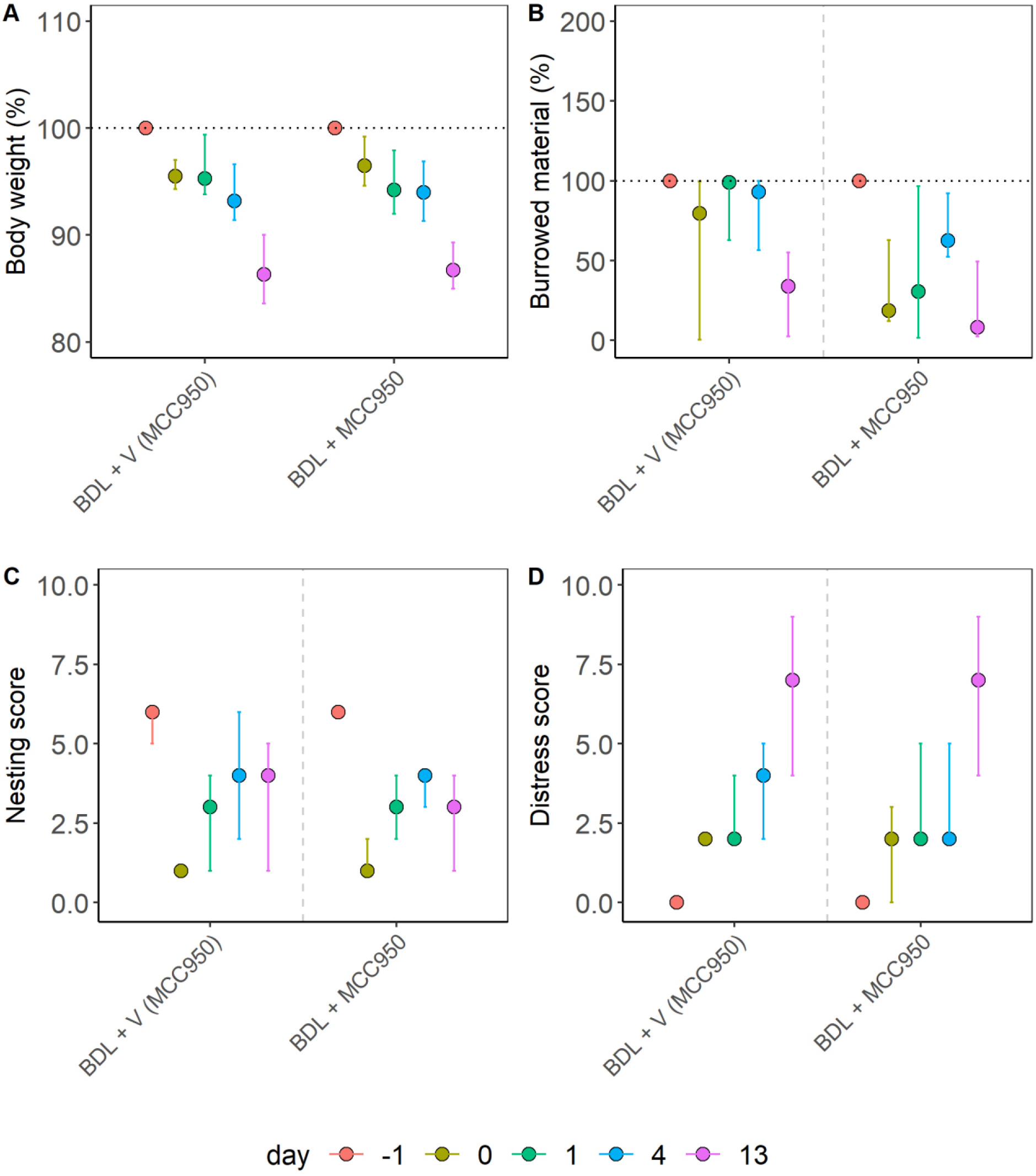
Distress evaluation of a cholestasis model in mice. Cholestasis was induced by bile duct ligation (BDL), and the animals were treated with an NLRP3 inflammasome inhibitor (MCC950) or the appropriate vehicle solution (V). The percentage of body weight (A), the percentage of burrowing activity (B), the nesting (C), and the distress score (D) were evaluated on the indicated days. No significant differences between the groups were determined using the Kruskal-Wallis test in (A) (χ^2^= 0.16, df = 1, p = 0.69), B (χ^2^= 2.58, df = 1, p = 0.11), C (χ^2^= 0.05, df = 1, p = 0.82) or D (χ^2^= 0.26, df = 1, p = 0.61). The graphs depict the bootstrapped median estimator on each experimental day and the 95 % confidence intervals.

The distress of animal models and experimental subgroups (Figures 5-6) was calculated with the Relative Severity Assessment (RELSA) algorithm [45] from the RELSA R-package (https://talbotsr.com/RELSA/index.html). The RELSA score was determined using three input variables (body weight, burrowing activity, and distress score) mapped against a reference set (laboratory A) with the same variables but from an independent transmitter-implantation experiment with a defined qualitative severity (i.e., moderate severity in laboratory B). Further, the highest RELSA_max_ value was obtained for each individual as the maximum RELSA during the treatment, representing the most experienced quantitative distress. The resulting RELSA_max_ values in the experimental subgroups were 10000-fold bootstrapped to obtain estimates on the median 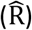 and the corresponding 95 % confidence intervals. Between-subgroup/model comparisons were calculated with the Wilcoxon-Mann-Whitney test in case of evidence for non-normal data, followed by Holm’s correction to determine differences in distress levels. In the case of normally distributed data, the adjusted *t*-test was used. The robustness of the severity estimation was tested with data from a second laboratory (B), using only body weight and burrowing activity as input variables. RELSA_max_ values >1 were considered more severe than the reference model.

**Fig. 5:**
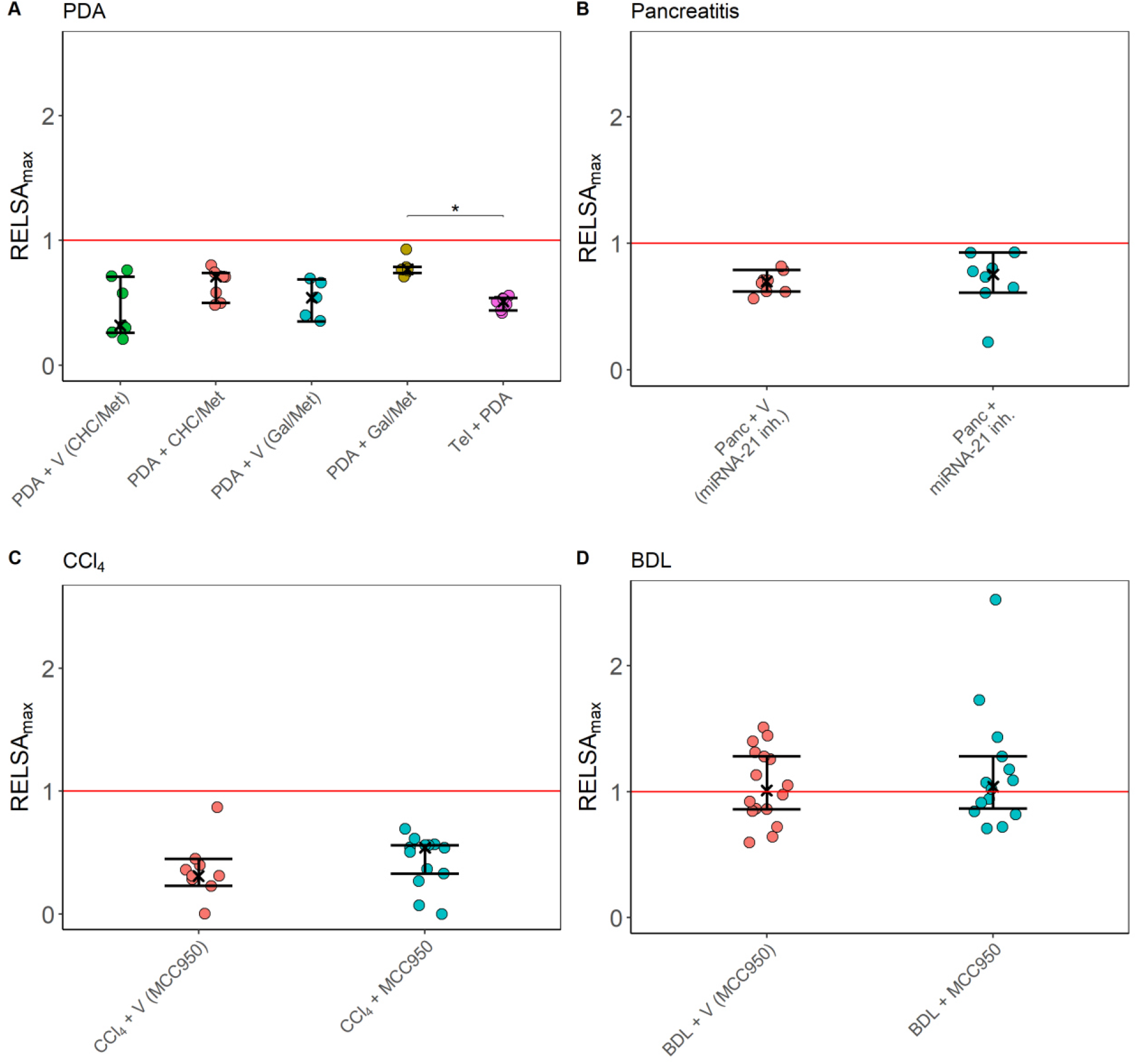
*Within*-model comparisons of distress using the variables body weight, burrowing activity, and the distress score, represented as the RELSA_max_ metric. The red line denotes the maximally experienced severity in the telemetry experiment of laboratory A (reference line), also on the RELSA scale. In each animal model, the distress was assessed between treatment groups. The panels show PDA (A), pancreatitis (B), carbon tetrachloride (CCl_4_) (C), and the cholestasis (BDL) model (D) – always in comparison to the reference level. Distinct treatments with drugs (CHC/Met, Gal/Met, miRNA-21 inh., MCC950) or treatment with appropriate vehicle controls (V) are indicated. The 95 % confidence intervals in all treatment groups remain below the reference line, indicating no evidence that any analyzed treatment has a higher severity than the surgery model. The Kruskal-Wallis test revealed a significant difference between the treatment groups only in panel A (χ^2^= 16.12, df = 4, p-value = 0.003), more specifically between PDA + Gal/Met and Tel + PDA (*p_adj_= 0.021). The graphs depict the RELSA_max_ values obtained from individual RELSA analyses, the bootstrapped median estimators, and the 95 % confidence intervals.

**Fig. 6:**
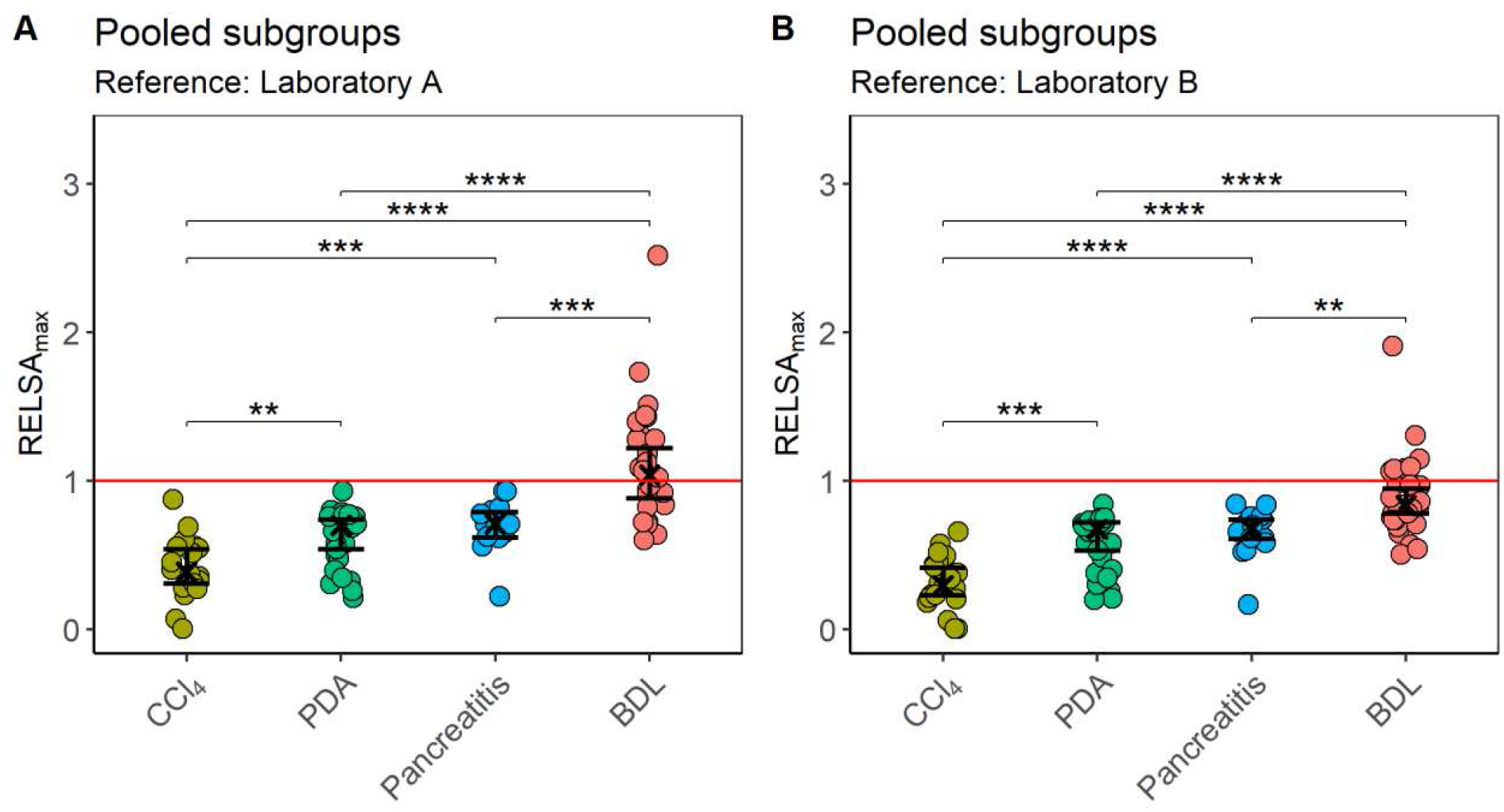
*Between*-model comparisons of distress after pooling the experimental subgroups. (A) The distress of the carbon tetrachloride (CCl_4_), pancreatitis (Panc), pancreatic cancer (PDA), and the cholestasis (BDL) model was evaluated using the RELSA_max_ values, with the reference built from data of laboratory A (transmitter implantation model, measured variables: body weight, burrowing activity as well as the distress score, the red line denotes the maximally experienced severity in the telemetry experiment). A Kruskal-Wallis test shows significant differences between the RELSA_max_ estimates of the animal models in panel A (χ^2^= 54.86, df = 3, p<0.0001). Differences were observed between CCl_4_ and PDA (**p_adj_ <0.01), CCl_4_ and BDL (****p_adj_ <0.0001), pancreatitis and BDL (***p_adj_ <0.001) as well as between PDA and BDL (****p_adj_ <0.0001). The following order of model severity can be seen by ordering the RELSA_max_ estimates: 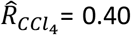, CI_95%_[0.29; 0.52]) < PDA (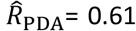, CI_95%_[0.50; 0.71]) < Pancreatitis (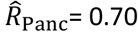, CI_95%_[0.56; 0.83]) < BDL (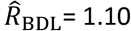, CI_95%_[1.00; 1.20]). (B) Testing the robustness of the procedure by implementing a different reference set from laboratory B with the variables body weight and burrowing activity. The treatment groups also showed significant differences using the Kruskal-Wallis test (χ^2^= 53.15, df = 3, p<0.0001). Pairwise differences were observed between CCl_4_ and PDA (***p_adj_ <0.001), CCl_4_ and Pancreatitis (****p_adj_<0.0001), CCl_4_ and BDL (****p_adj_ <0.0001), Pancreatitis and BDL (**p_adj_ <0.01) as well as between PDA and BDL (****p_adj_ <0.0001). Here, the order of RELSA_max_ estimates showed as follows: 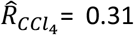, CI_95%_[0.22; 0.40]) < PDA (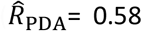, CI_95%_[0.50; 0.67]) < Pancreatitis (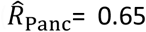, CI_95%_[0.54; 0.76]) < BDL (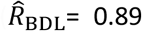, CI_95%_[0.81; 0.96]). The graphs depict the RELSA_max_ values obtained from individual RELSA analyses (pooled subgroups for each animal model) and the bootstrapped median estimator together with the 95 % confidence intervals.

### Data availability

Raw data can be downloaded from a GitHub repository under the following link: https://github.com/mytalbot/gastrointestinal_data.

## Results

### Evaluation of single readout parameters for distress

In particular animal models, we analyzed four readout parameters for animal distress (body weight change, burrowing activity, nesting behavior, and a distress score) to assess the robustness of distress evaluation. Within each animal model, distinct groups of mice experienced slight variations in the experimental procedures. For example, the animals were treated with various drugs or vehicle solutions or had an implanted telemetry transmitter.

When assessing an animal model for pancreatic ductal adenocarcinoma (PDA), no significant differences in body weight change were observed when comparing groups of mice treated with specific drugs or vehicle solutions (Fig. 1A). However, significant differences in body weight change were observed between mice, which had a telemeter implanted and telemeter-free mice treated with specific drug combinations (Fig. 1A). No significant differences were also observed when analyzing burrowing activity and nesting behavior. Still, a significant difference was observed in the distress score when comparing distinct groups of mice (Fig. 1B-D). We concluded that some readout parameters, such as burrowing activity, are very robust to slight variations in the experimental procedure. In contrast, other readout parameters, such as the distress score, are prone to experimental variation.

We also evaluated identical readout parameters for distress in an animal model for chronic pancreatitis and compared the distress between groups of mice treated either with a drug or vehicle control (Fig. 2). No significant differences between these two groups were detected when analyzing body weight change, burrowing activity, nesting behavior, or a distress score (Fig. 2).

In addition, we assessed two animal models for liver damage. Liver damage was either caused by redundant carbon tetrachloride (CCl_4_) administration or by cholestasis induced by bile duct ligation (BDL). When analyzing the CCl_4_ animal model, no significant differences between treatments were detected (Fig. 3). Also, no significant differences between treatments were found in the BDL model (Fig. 4).

These data demonstrated that the distress readout parameters in three out of four animal models were robust to slight variations in the experimental procedure (Fig. 2-4). In contrast, body weight and distress score gave different results in the PDA animal model when the experimental procedures varied slightly (Fig. 1). Since we noticed that other readout parameters for distress gave varying results (Fig. 1), we developed a method that combined multiple variables throughout a treatment into a singular score, the maximum of the Relative Severity Assessment score (RELSA_max_) [45].

### Combining multiple variables into RELSA scores

RELSA is a weighted procedure that maps multiple input variables into a single metric, allowing the comparison of different animal models and measurements on a quantitative scale. The comparability between models was achieved by referencing the standardized data to a set with a well-defined qualitative and quantitative severity. In the reference data, three variables from an intraperitoneal transmitter implantation experiment were used (body weight change, burrowing activity, and the distress score, Fig. S1, laboratory A). The measurements showed that the loss in body weight, burrowing activity, and distress score was largest on day 0 (the first day after surgery). Over time, the values recovered back to baseline levels. According to the RELSA concept, the experimental measurements were compared against the maximum deviations in the reference set, scaled, and combined into the RELSA score. The RELSA procedure gives more attention to larger deviations so that potential noise gets less weight. The maximum RELSA value per treatment (RELSA_max_) was then used to compare models on a time-independent basis.

### *Within*-model comparisons of distress using RELSA_max_

The bootstrapped RELSA_max_ estimates of the five pancreatic cancer experimental subgroups (Fig. 5A) were all below the RELSA reference level of laboratory A. Also, the 95 % confidence intervals of the subgroup estimates did not cross the reference line. Therefore, it can be concluded that pancreatic cancer models show significantly lower distress than the telemetric implantation model. The between-subgroup comparisons showed that the highest distress was achieved in the PDA + Gal/Met subgroup. In addition, the RELSA_max_ of these animals was significantly higher than those within the Tel + PDA subgroup (Fig. 5A, p_adj_=0.021).

In the chronic pancreatitis model (Fig. 5B), the 95 % confidence intervals of the bootstrapped RELSA_max_ estimates in the two subgroups were lower than the reference set, indicating significantly lower distress than in the telemetric implantation model. No significant difference between these subgroups was detected

In the two CCl_4_ subgroups, the 95 % confidence intervals of the bootstrapped RELSA_max_ estimates were also lower than the reference set, indicating significantly lower distress than in the telemetric implantation model (Fig. 5C). Also, in this animal model, no significant difference between the subgroups was detected (Fig. 5C).

In the BDL model, both estimates were above the reference level (Fig. 5D). However, the lower confidence intervals remained below the reference level. Therefore, there was insufficient evidence to support the hypothesis that the BDL model leads to more severe distress than the reference model. However, two animals in the BDL + MCC950 subgroup experienced very high distress (e.g., RELSA_max_=1.73 and RELSA_max_=2.52). Both animals reached the humane endpoint (> 20 % body weight reduction) at the end of the experiment. Again, no significant difference between the subgroups of this animal model was found.

### *Between*-model comparisons of distress using RELSA_max_

The distinct subgroups were pooled to focus more on comprehensive model comparisons than experiments. Again, the RELSA_max_ procedure was used to compare distress levels between four animal models (Fig. 6A). This was achieved by mapping the experimental data to the standardized severity of the reference data from laboratory A. The order of estimates yielded a ranking of severity used to classify the animal models in terms of distress magnitude. This ranking showed the following order in increasing severity based on the RELSA_max_ model estimates: CCl_4_ < PDA ≈ Pancreatitis < BDL. Significant differences were found when comparing these animal models (Fig. 6A). Pancreatitis showed significantly higher severity than the CCl_4_ model (p_adj_≤0.001), which was also true for CCl_4_ vs. PDA (p_adj_=0.01). Further, the BDL model showed significantly higher severity towards the CCl_4_ (p_adj_≤0.0001), pancreatitis (p_adj_≤0.001), and PDA (p_adj_≤0.0001) models.

The robustness of this severity ranking was tested with an independent reference set using two variables (body weight change and burrowing activity) from an intraperitoneal transmitter implantation experiment in laboratory B. Both variables showed the same pattern of estimates on day 0 with subsequent recovery during the following days (see raw data of laboratory A Fig. S1 and laboratory B Fig. S2).

Testing the four animal models against the reference data from laboratory B resulted in RELSA_max_ estimates that were slightly lower than the ones from the first analysis (see Fig. 6A and B). However, this validation analysis still maintained the previous order of model severity: CCl_4_ < PDA ≈ Pancreatitis < BDL (Fig. 6B). Again, pancreatitis (p_adj_≤0.0001) and PDA (p_adj_≤0.001) showed significantly higher severity than the CCl_4_ model. Further, the BDL model showed significantly higher severity towards the CCl_4_ (p_adj_≤0.0001), pancreatitis (p_adj_≤0.01), and PDA (p_adj_≤0.0001) models. Thus, identical significant differences were observed between the four animal models when using both reference sets. Therefore, judging the severity of animal models based on the significant differences in distress using the RELSA_max_ method was robust towards using different reference data.

## Discussion

This study evaluated animal distress by assessing the body weight change, a distress score, and the burrowing and nesting behavior in four animal models for distinct gastrointestinal diseases (Fig. 1-4). The robustness within each model was evaluated by minor experimental design modifications, e.g., different treatment strategies. The previously established RELSA procedure [45] graded the maximum distress each animal experiences by mapping the multi-dimensional information of various distress parameters against a specific reference set of defined severity. The RELSA_max_ score proved robust in three out of four animal models when comparing each animal model under different experimental conditions (Fig. 5). The robustness was also given when using RELSA reference sets from two laboratories to estimate the order of severity in the analyzed animal models with the RELSA_max_ value (Fig. 6).

The basis for an evidence-based severity assessment of animal models is the use of reliable distress parameters [25,26,55]. A robust body weight reduction was observed in two out of four animal models (BDL and pancreatitis) as a response to surgical intervention or chemical induction (Fig. 1-4), which implies this parameter’s relevance to detecting distress in some gastrointestinal animal models. Body weight alone or combined with other criteria is useful for humane endpoint determination in many animal models [34,56–58]. Its evaluation is highly recommended for severity grading by welfare assessment protocols [25,26]. A robust reduction of burrowing activity was observed in the BDL, pancreatitis, and PDA animal model, but not after CCl_4_ intoxication (thus, robustness was given in three out of four animal models). A robust reduction of nesting activity was also observed in three out of four animal models (BDL, CCl_4_, and pancreatitis). The results indicated that burrowing and nesting behavior were even more robust in these experiments than in body weight reduction. Indeed, both behavior tests have been reported to detect stress and suffering in many distinct animal models. For example, in animal models for colitis [59], Parkinson disease [60], pancreatitis [61], epilepsy [62,63], and depression [64]. However, a robust increase in the distress score could only be detected in the BDL animal model. This result indicates that in some animal models, the distress score is less informative than body weight change or behavior tests. This conclusion is consistent with previous studies [47].

However, we also want to describe specific limitations to the abovementioned summary and conclusions. For example, within the PDA model, significant differences in body weight change were detected between the Tel + PDA and PDA + CHC/Met or PDA + Gal/Met group (Fig. 1A). An initial body weight reduction caused these differences as a result of the previous surgery for transmitter implantation and no additional decrease of body weight in the Tel + PDA group after cell injection [46]. Thus, body weight reduction is robust if one only compares PDA animal models using different pharmacological treatments. Still, it is not robust when an entire additional surgical intervention (telemeter implantation) is included in the PDA model. Furthermore, low robustness was also observed when comparing the distress score between PDA + GAL/Met and other treatment groups. This result most likely reflects the side effects of specific drugs, as described in a previous study [47]. Thus, this is an example that the robustness of data within an animal model might be reduced if the changed variable, e.g., therapy, causes an intense effect on the well-being of the animals.

We observed low robustness of methods when only single parameters were used to compare the distress between animal models. For example, the BDL model causes a gradual increase in the distress score due to a continuous progression of the disease (Fig. 4D). In contrast, after CCl_4_ injection, no increase in the distress score could be observed during chronic pancreatitis (Fig. 2D). Possibly, methods measure distress in an animal-model specific manner. This outcome was also marked by Mallien et al. when genetic, stress-based, and pharmacological mouse models of depression were compared [64]. These differences between animal models highlight the need to perform multi-parametric severity assessment when comparing different animal models [45,55].

In the present study, a multi-parametric animal model assessment was done using the RELSA algorithm. This algorithm allowed an informed integration of various experimentally available read-our parameters into a single value. The maximum deviations per animal observed on the RELSA scale during an experiment were named RELSA_max_. These values represent the utmost distress animals experience on a quantitative scale compared to a defined reference set [45]. Since it is recommended by the EU Commission to consider the highest distress an animal experiences for defining the severity grade of an experiment [65], the RELSA_max_ is an excellent tool to determine and compare severity levels between animal models.

Please note that variables with large weights will contribute more information to the final RELSA score than variables with little weight. Thus, specific markers may dominate the RELSA values due to their model-specificity. However, these values are adequately mapped to the RELSA space due to the weighting system of the algorithm. Therefore, a holistic comparison of animal models and individual animals is possible with the limitation that the reference set must at least partially contain the same measured variables as the tested models.

No significant differences in the RELSA_max_ were observed, within each animal model, when comparing their varying experimental conditions, except when analyzing the PDA model (Fig. 5). Here, the significant difference between PDA + Gal/Met and Tel + PDA model can be explained by the lack of body weight reduction in the Tel + PDA model and a high distress score in the PDA + Gal/Met group (for a detailed discussion see previous text).

When pooling the data from various experiments for each animal model and comparing the severity of all animal models with the RELSA_max_ (Fig. 6), the CCl_4_, PDA, and pancreatitis models were below a RELSA_max_ of 1.0. This result indicates that these animal models cause less distress than transmitter implantation, which was used as a reference. Interestingly, the RELSA_max_ not only allows a comparison of the severity of various animal models to the reference model but is also an excellent tool to compare the distress between different animal models (Fig. 6). Based on RELSA_max_, the CCl_4_ model indicates the lowest severity. In contrast, the PDA model is significantly more stressful for the animals. The pancreatitis model is shown to have a similar severity level to the PDA model. The BDL model has the highest severity, indicated by a significantly higher RELSA_max_ than the other gastrointestinal animal models. The progression of cholestasis leads to a substantial impairment of welfare, as characterized by major changes in all four single distress parameters (Fig. 4). That this animal model is quite severe is also supported by the low survival rate ranging from 64-70 % [66–68]. In contrast, the survival rates of pancreatitis at 99 % [61], the CCl_4_ model at 100 % [69], and the PDA model at 83 % [70] were reported to be higher.

Suppose one considers the implantation of a transmitter as a surgical intervention, which causes moderate distress as suggested by the EU-Commission Guideline in Annex XIII [11]. In that case, one can define the distress caused by other animal models by implementing the RELSA procedure. Based on this concept, the CCl_4,_ PDA, and pancreatitis models might cause mild to moderate distress, whereas the BDL model might cause moderate to severe distress. However, please note that we compare only the highest distress level, which an animal reaches during the experiment at a single time point. When analyzing the RELSA_max_ values, we do not consider how long an animal is distressed during an experiment. Thus, cumulative suffering might have to be considered in addition to the RELSA_max_ when defining specific severity categories. Such longitudinal data can be presented using standard RELSA curves [45]. However, a concept of how to summarize cumulative suffering by an algorithm to allow direct comparison between, for example, short-term severe distress to long-term moderate distress still needs to be developed.

## Conclusion

The present study characterized the robustness of distress assessment using multiple non-invasive methods. With the implementation of the RELSA procedure, the highest distress levels in animals during experiments were described mathematically with the RELSA_max_ value. This score allowed us to perform several *between*-animal model comparisons and reference tests. Since the results were very robust, the RELSA_max_ is reliable for comparing animal models. The algorithm might also be valuable when examining drug side effects or evaluating refinement measures during *in vivo* experiments.

## Figure legends

**Supplementary Fig. 1:** Description of the reference set (implantation of a telemetric transmitter in laboratory A). The percentage of body weight (A), the percentage of burrowing activity (B), and a distress score (C) were evaluated on the indicated days. (A) The daily data showed normally distributed estimators after the bootstrapping in the percent body weight variable. Therefore, a *repeated-measures* ANOVA was used to determine differences between experimental days. The percent body weight variable showed significant differences over time (F(6,54)=56.562, p<0.0001, η_G_^2^=0.672). Dunnett’s post hoc test was used to determine significant differences on the indicated days compared to day -1 (**p_adj_≤0.01, ***p_adj_≤0.001, ****p_adj_≤0.0001). (B) The burrowing data did not show normal distribution on the experimental days. Therefore, a Friedman test with the day as the *within-subjects* variable was performed to assess the development over time. The burrowing variable showed significant differences over time (χ^2^=33.9, df=6, p<0.0001). Dunn’s post-hoc test with Holm’s correction revealed that there were significant differences between day -1 (baseline) and day 0 (p_adj_<0.0001). (C) The Friedman test of the distress score showed a significant difference between days as the *within-subjects* variable (χ^2^=48.6, df=6, p<0.0001). Dunn’s post-hoc test with Holm’s correction shows a significant difference between day -1 (control) and day 0 (p_adj_<0.0001). The graphs depict the bootstrapped median estimator on each experimental day and the 95 % confidence intervals.

**Supplementary Fig. 2:** Description of the validation reference set (implantation of a telemetric transmitter in laboratory B). The percentage of body weight (A) and the percentage of burrowed material (B) were evaluated on the indicated days. (A) The daily data showed normally distributed estimators after the bootstrapping in the percent body weight variable. Therefore, a *repeated-measures* ANOVA was calculated to determine differences between experimental days. The percent body weight variable showed significant differences over time (F(5,60)=82.01, p<0.0001, η_G_^2^=0.746). Dunnett’s post-hoc test was used to determine significant differences (***p_adj_≤0.001) at the indicated days compared to baseline (day -1). (B) The burrowing data did not show normal distribution on the experimental days. Therefore, a Friedman test with the day as the *within*-subjects variable was performed to assess the development over time. The burrowing variable showed significant differences over time (χ^2^=43.874, df=5, p<0.0001). Dunn’s post-hoc test with Holm’s correction revealed that there was a significant difference between day -1 and day 0 (****p_adj_<0.0001) as well as between days -1 and 1 (*p_adj_=0.026). The graphs depict the bootstrapped median estimator on each experimental day and the 95 % confidence intervals.

## Acknowledgments

This study was supported by the Deutsche Forschungsgemeinschaft (DFG research group FOR 2591, ZE 712/1-1, ZE 712/1-2, VO 450/15-1, and VO 450/15-2 as well as HA6483/1-2, BL953/10-1 and 10-2, BL953/11-1 and 11-2). In addition, Guanglin Tang was supported by the China Scholarship Council (grant number: 201808080167).

## References

1. Sims, E. K., Carr, A. L. J., Oram, R. A., DiMeglio, L. A. & Evans-Molina, C. 100 years of insulin: celebrating the past, present and future of diabetes therapy. Nat Med 27, 1154–1164. http://doi.org/10.1038/s41591-021-01418-2 (2021).

2. Barré-Sinoussi, F. & Montagutelli, X. Animal models are essential to biological research: issues and perspectives. Future Science OA 1. http://doi.org/10.4155/fso.15.63 (2015).

3. van Tilbeurgh, M. et al. Predictive Markers of Immunogenicity and Efficacy for Human Vaccines. Vaccines 9, 579. http://doi.org/10.3390/vaccines9060579 (2021).

4. Phillips, N. L. H. & Roth, T. L. Animal Models and Their Contribution to Our Understanding of the Relationship Between Environments, Epigenetic Modifications, and Behavior. Genes 10, 47. http://doi.org/10.3390/genes10010047 (2019).

5. Ohl, F. & van der Staay, F. J. Animal welfare: At the interface between science and society. The Veterinary Journal 192, 13–19. http://doi.org/10.1016/j.tvjl.2011.05.019 (2012).

6. Gross, D. & Tolba, R. H. Ethics in Animal-Based Research. Eur Surg Res 55, 43–57. http://doi.org/10.1159/000377721 (2015).

7. Petetta, F. & Ciccocioppo, R. Public perception of laboratory animal testing: Historical, philosophical, and ethical view. Addiction Biology. http://doi.org/10.1111/adb.12991 (2020).

8. Codecasa, E., Pageat, P., Marcet-Rius, M. & Cozzi, A. Legal Frameworks and Controls for the Protection of Research Animals: A Focus on the Animal Welfare Body with a French Case Study. Animals 11, 695. http://doi.org/10.3390/ani11030695 (2021).

9. Lee, K. H., Lee, D. W. & Kang, B. C. The ‘R’ principles in laboratory animal experiments. Lab Anim Res 36. http://doi.org/10.1186/s42826-020-00078-6 (2020).

10. Olsson, I. A. S., Silva, S. P. d., Townend, D. & Sandøe, P. Protecting Animals and Enabling Research in the European Union: An Overview of Development and Implementation of Directive 2010/63/EU. ILAR Journal 57, 347–357. http://doi.org/10.1093/ilar/ilw029 (2017).

11. European Parliament. Directive 2010/63/EU of the European Parliament and of the Council of 22 September 2010 on the protection of animals used for scientific purposesText with EEA relevance (2010).

12. United States Department of Agriculture. USDA Animal Care: Animal Welfare Act and Animal Welfare Regulations. Available at https://www.aphis.usda.gov/animal_welfare/downloads/bluebook-ac-awa.pdf (2019).

13. MacArthur Clark, J. A. & Sun, D. Guidelines for the ethical review of laboratory animal welfare People’s Republic of China National Standard GB/T 35892-2018 [Issued 6 February 2018 Effective from 1 September 2018]. Animal Model Exp Med 3, 103–113. http://doi.org/10.1002/ame2.12111 (2020).

14. Foley, P. L., Kendall, L. V. & Turner, P. V. Clinical Management of Pain in Rodents. comp med 69, 468–489. http://doi.org/10.30802/AALAS-CM-19-000048 (2019).

15. Herrmann, K. & Flecknell, P. The Application of Humane Endpoints and Humane Killing Methods in Animal Research Proposals: A Retrospective Review. Altern Lab Anim 46, 317–333. http://doi.org/10.1177/026119291804600606 (2018).

16. Keubler, L. M. et al. Where are we heading? Challenges in evidence-based severity assessment. Lab Anim 54, 50–62. http://doi.org/10.1177/0023677219877216 (2020).

17. Bleich, A. & Tolba, R. H. How can we assess their suffering? German research consortium aims at defining a severity assessment framework for laboratory animals. Lab Anim 51, 667. http://doi.org/10.1177/0023677217733010 (2017).

18. Talbot, S. R. et al. Defining body-weight reduction as a humane endpoint: a critical appraisal. Lab Anim 54, 99–110. http://doi.org/10.1177/0023677219883319 (2020).

19. Morton, D. B. & Griffiths, P. H. Guidelines on the recognition of pain, distress and discomfort in experimental animals and an hypothesis for assessment. The Veterinary record 116, 431–436. http://doi.org/10.1136/vr.116.16.431 (1985).

20. Paster, E. V., Villines, K. A. & Hickman, D. L. Endpoints for mouse abdominal tumor models: refinement of current criteria. comp med 59, 234–241 (2009).

21. Deacon, R. M. J. Burrowing in rodents: a sensitive method for detecting behavioral dysfunction. Nat Protoc 1, 118–121. http://doi.org/10.1038/nprot.2006.19 (2006).

22. Gjendal, K., Ottesen, J. L., Olsson, I. A. S. & Sørensen, D. B. Burrowing and nest building activity in mice after exposure to grid floor, isoflurane or ip injections. Physiology & Behavior 206, 59–66. http://doi.org/10.1016/j.physbeh.2019.02.022 (2019).

23. Kahnau, P., Habedank, A., Diederich, K. & Lewejohann, L. Behavioral Methods for Severity Assessment. Animals 10, 1136. http://doi.org/10.3390/ani10071136 (2020).

24. Zechner, D. et al. Generalizability, Robustness and Replicability When Evaluating Wellbeing of Laboratory Mice with Various Methods. Animals 12, 2927. http://doi.org/10.3390/ani12212927 (2022).

25. Smith, D. et al. Classification and reporting of severity experienced by animals used in scientific procedures: FELASA/ECLAM/ESLAV Working Group report. Lab Anim 52, 5–57. http://doi.org/10.1177/0023677217744587 (2018).

26. Hawkins, P. et al. A guide to defining and implementing protocols for the welfare assessment of laboratory animals: eleventh report of the BVAAWF/FRAME/RSPCA/UFAW Joint Working Group on Refinement. Lab Anim 45, 1–13. http://doi.org/10.1258/la.2010.010031 (2011).

27. Harikrishnan, V. S., Hansen, A. K., Abelson, K. S. P. & Sørensen, D. B. A comparison of various methods of blood sampling in mice and rats: Effects on animal welfare. Lab Anim 52, 253–264. http://doi.org/10.1177/0023677217741332 (2018).

28. Hurst, J. L. & West, R. S. Taming anxiety in laboratory mice. Nat Methods 7, 825–826. http://doi.org/10.1038/nmeth.1500 (2010).

29. Lofgren, J. et al. Analgesics promote welfare and sustain tumour growth in orthotopic 4T1 and B16 mouse cancer models. Lab Anim 52, 351–364. http://doi.org/10.1177/0023677217739934 (2018).

30. Peng, M. et al. Battery of behavioral tests in mice to study postoperative delirium. Sci Rep 6. http://doi.org/10.1038/srep29874 (2016).

31. Ebrahimi Kalan, M., Jebai, R., Zarafshan, E. & Bursac, Z. Distinction Between Two Statistical Terms: Multivariable and Multivariate Logistic Regression. Nicotine & tobacco research : official journal of the Society for Research on Nicotine and Tobacco 23, 1446–1447. http://doi.org/10.1093/ntr/ntaa055 (2021).

32. Wassermann, L. et al. Monitoring of Heart Rate and Activity Using Telemetry Allows Grading of Experimental Procedures Used in Neuroscientific Rat Models. Frontiers in neuroscience 14, 587760. http://doi.org/10.3389/fnins.2020.587760 (2020).

33. Häger, C. et al. Running in the wheel: Defining individual severity levels in mice. PLoS biology 16, e2006159. http://doi.org/10.1371/journal.pbio.2006159 (2018).

34. Helgers, S. O. A. et al. Body weight algorithm predicts humane endpoint in an intracranial rat glioma model. Sci Rep 10, 9020. http://doi.org/10.1038/s41598-020-65783-7 (2020).

35. Bruch, S., Ernst, L., Schulz, M., Zieglowski, L. & Tolba, R. H. Best variable identification by means of data-mining and cooperative game theory. Journal of biomedical informatics 113, 103625. http://doi.org/10.1016/j.jbi.2020.103625 (2021).

36. Efron, B. Bootstrap Methods: Another Look at the Jackknife. Ann. Statist. 7. http://doi.org/10.1214/aos/1176344552 (1979).

37. Narİnç, D., Aygün, A., KüçüköNder, H., Aksoy, T. & GüRCAN, E. K. Hayvancılık Alanında Bootstrap Tekniğinin Bir Uygulaması: Yumurta Sarı Rengi Örneği. Kafkas Univ Vet Fak Derg. http://doi.org/10.9775/kvfd.2014.12693 (2015).

38. Wood, M. Statistical inference using bootstrap confidence intervals. Significance 1, 180–182. http://doi.org/10.1111/j.1740-9713.2004.00067.x (2004).

39. Lee, D. K. Alternatives to P value: confidence interval and effect size. Korean journal of anesthesiology 69, 555–562. http://doi.org/10.4097/kjae.2016.69.6.555 (2016).

40. Sim, J. & Reid, N. Statistical Inference by Confidence Intervals: Issues of Interpretation and Utilization. Physical Therapy 79, 186–195. http://doi.org/10.1093/ptj/79.2.186 (1999).

41. Goodman, S. N., Fanelli, D. & Ioannidis, J. P. A. What does research reproducibility mean? Sci. Transl. Med. 8, 341ps12–341ps12. http://doi.org/10.1126/scitranslmed.aaf5027 (2016).

42. Erdogan, B. R. & Michel, M. C. in Good research practice in non-clinical pharmacology and biomedicine, edited by A. Bespalov, M. C. Michel & T. Steckler (Springer Open, 2020), pp. 163– 175.

43. Pallocca, G., Rovida, C. & Leist, M. On the usefulness of animals as a model system (part I): Overview of criteria and focus on robustness. ALTEX 39, 347–353. http://doi.org/10.14573/altex.2203291 (2022).

44. Strech, D. & Dirnagl, U. 3Rs missing: animal research without scientific value is unethical. BMJ Open Science 3, bmjos-2018-000048. http://doi.org/10.1136/bmjos-2018-000048 (2019).

45. Talbot, S. R. et al. RELSA—A multidimensional procedure for the comparative assessment of well-being and the quantitative determination of severity in experimental procedures. Front. Vet. Sci. 9. http://doi.org/10.3389/fvets.2022.937711 (2022).

46. Kumstel, S. et al. Benefits of non-invasive methods compared to telemetry for distress analysis in a murine model of pancreatic cancer. Journal of advanced research 21, 35–47. http://doi.org/10.1016/j.jare.2019.09.002 (2020).

47. Kumstel, S. et al. Grading animal distress and side effects of therapies. Annals of the New York Academy of Sciences 1473, 20–34. http://doi.org/10.1111/nyas.14338 (2020).

48. Abdelrahman, A. et al. A novel multi-parametric analysis of non-invasive methods to assess animal distress during chronic pancreatitis. Sci Rep 9, 14084. http://doi.org/10.1038/s41598-019-50682-3 (2019).

49. Tang, G. et al. Comparing distress of mouse models for liver damage. Sci Rep 10, 19814. http://doi.org/10.1038/s41598-020-76391-w (2020).

50. Deacon, R. Assessing burrowing, nest construction, and hoarding in mice. Journal of visualized experiments : JoVE, e2607. http://doi.org/10.3791/2607 (2012).

51. Kumstel, S. et al. Grading Distress of Different Animal Models for Gastrointestinal Diseases Based on Plasma Corticosterone Kinetics. Animals : an open access journal from MDPI 9. http://doi.org/10.3390/ani9040145 (2019).

52. R Core Team. R: A language and environment for statistical computing. R Foundation for Statistical Computing, Vienna, Austria. Available at https://www.R-project.org/. (2020).

53. Mangiafico, S. rcompanion: Functions to Support Extension Education Program Evaluation. R package version 2.3.27. Available at http://rcompanion.org/ (2021).

54. Kassambara, A. rstatix: Pipe-Friendly Framework for Basic Statistical Tests. Available at https://github.com/kassambara/rstatix (2021).

55. Keubler, L. M. et al. Where are we heading? Challenges in evidence-based severity assessment. Lab Anim 54, 50–62. http://doi.org/10.1177/0023677219877216 (2020).

56. How to determine humane endpoints for research animals. Lab animal 45, 19. http://doi.org/10.1038/laban.908 (2016).

57. Hankenson, F. C. et al. Weight loss and reduced body temperature determine humane endpoints in a mouse model of ocular herpesvirus infection. j am assoc lab anim sci 52, 277–285 (2013).

58. Mei, J. et al. Refining humane endpoints in mouse models of disease by systematic review and machine learning-based endpoint definition. ALTEX 36, 555–571. http://doi.org/10.14573/altex.1812231 (2019).

59. Cheatham, S. M. et al. Morphine Exacerbates Experimental Colitis-Induced Depression of Nesting in Mice. Frontiers in pain research (Lausanne, Switzerland) 2, 738499. http://doi.org/10.3389/fpain.2021.738499 (2021).

60. Sager, T. N. et al. Nest building performance following MPTP toxicity in mice. Behavioural brain research 208, 444–449. http://doi.org/10.1016/j.bbr.2009.12.014 (2010).

61. Durst, M. et al. Analysis of Pain and Analgesia Protocols in Acute Cerulein-Induced Pancreatitis in Male C57BL/6 Mice. Front. Physiol. 12, 744638. http://doi.org/10.3389/fphys.2021.744638 (2021).

62. Boldt, L. et al. Toward evidence-based severity assessment in mouse models with repeated seizures: I. Electrical kindling. Epilepsy & behavior : E&B 115, 107689. http://doi.org/10.1016/j.yebeh.2020.107689 (2021).

63. van Dijk, R. M. et al. Design of composite measure schemes for comparative severity assessment in animal-based neuroscience research: A case study focussed on rat epilepsy models. PLoS ONE 15, e0230141. http://doi.org/10.1371/journal.pone.0230141 (2020).

64. Mallien, A. S. et al. Comparative Severity Assessment of Genetic, Stress-Based, and Pharmacological Mouse Models of Depression. Frontiers in behavioral neuroscience 16, 908366. http://doi.org/10.3389/fnbeh.2022.908366 (2022).

65. European Commission. Caring for animals aiming for better science. Severity Assessment framework. Available at https://ec.europa.eu/environment/chemicals/lab_animals/pdf/guidance/severity/en.pdf (2012).

66. Sigal, M. et al. Darbepoetin-α inhibits the perpetuation of necro-inflammation and delays the progression of cholestatic fibrosis in mice. Laboratory investigation; a journal of technical methods and pathology 90, 1447–1456. http://doi.org/10.1038/labinvest.2010.115 (2010).

67. Gäbele, E. et al. TNFalpha is required for cholestasis-induced liver fibrosis in the mouse. Biochemical and biophysical research communications 378, 348–353. http://doi.org/10.1016/j.bbrc.2008.10.155 (2009).

68. Zhang, X. et al. A rational approach of early humane endpoint determination in a murine model for cholestasis. ALTEX 37, 197–207. http://doi.org/10.14573/altex.1909111 (2020).

69. Duncan, M. B. et al. Type XVIII collagen is essential for survival during acute liver injury in mice. Disease models & mechanisms 6, 942–951. http://doi.org/10.1242/dmm.011577 (2013).

70. Kumstel, S. et al. Targeting pancreatic cancer with combinatorial treatment of CPI-613 and inhibitors of lactate metabolism. PLoS ONE 17, e0266601. http://doi.org/10.1371/journal.pone.0266601 (2022).

