## Supplemental Material for "Robustness of a multivariate composite score when evaluating distress of animal models for gastrointestinal diseases"

### Supplementary Figure S1

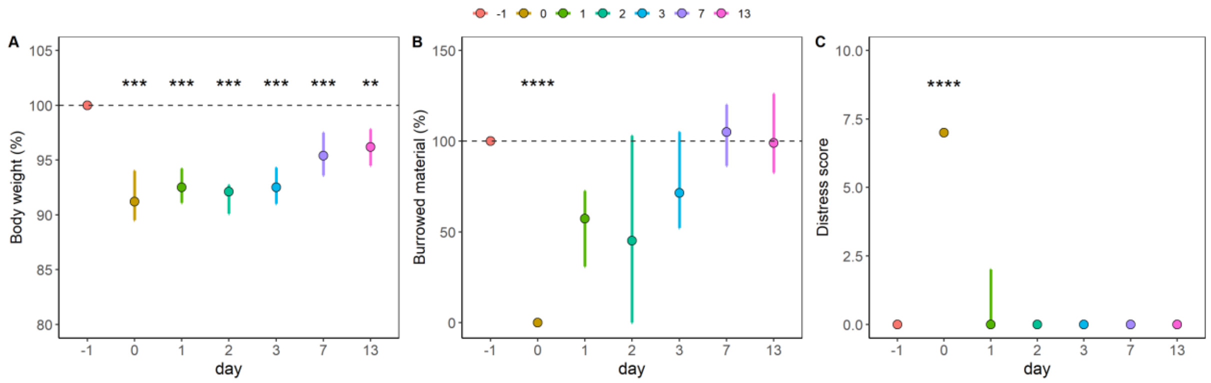


**S1.** Reference data laboratory A (surgery model, transmitter implantation) with (**A**) body weight change (%), (**B**) burrowed material (%), and (**C**) the distress score. Data are shown as bootstrapped estimates (median) with 95% confidence intervals. Significant baseline differences are indicated with *.

### Supplementary Figure S2

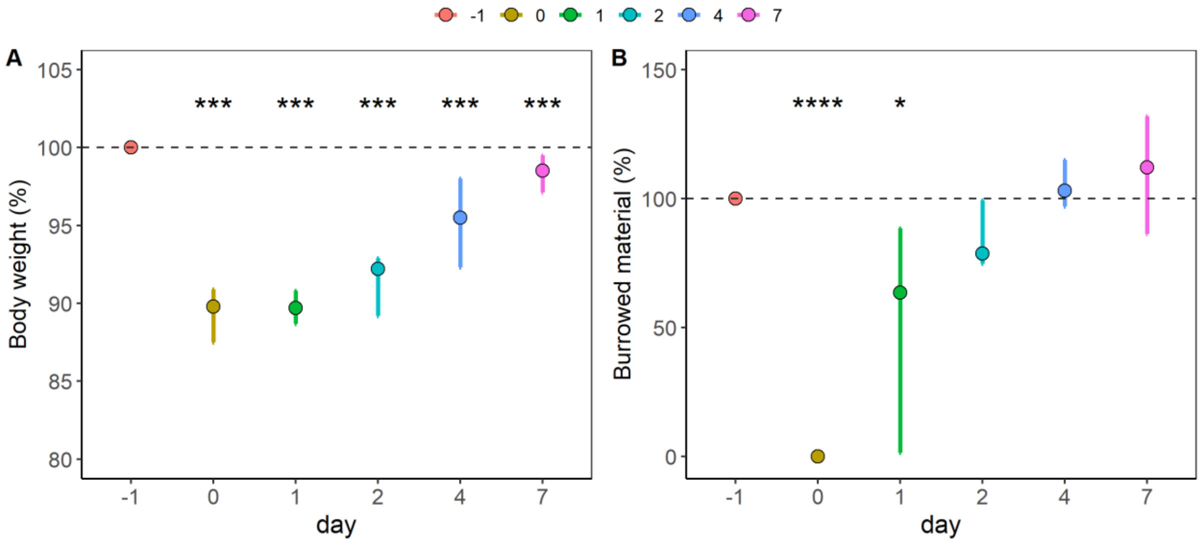


**S2.** Reference data laboratory B (surgery model, transmitter implantation) with (**A**) body weight change (%), (**B**) burrowed material (%). Note that the distress score is missing here. Data are shown as bootstrapped estimates (median) with 95% confidence intervals. Significant baseline differences are indicated with *.

### Supplementary Tables

The following tables (S1-S4) provide the p-values of the non-parametric Dunn's test for all pairwise comparisons of the experimental days vs. baseline values (day -1). The significant (Holm-corrected) *within*-subgroup comparisons are reported in the results as adjusted p-values ($p_{adj}$).

#### Table S1 - Pancreatic Cancer

**Body weight change.**

| day | PDA + V (CHC/Met) | PDA + CHC/Met | PDA + V (Gal/Met) | PDA + Gal/Met | Tel + PDA |
|---|---|---|---|---|---|
| **0** | 1.000 | 0.611 | 1.000 | 0.647 | 0.225 |
| **1** | 1.000 | 0.611 | 1.000 | 1.000 | 0.859 |
| **2** | 1.000 | 0.611 | 1.000 | 1.000 | 0.461 |
| **4-8** | 1.000 | 0.611 | 1.000 | 0.430 | 0.147 |
| **18-19** | 0.933 | 0.060 | 1.000 | 0.326 | 0.001 |
| **34-35** | 0.789 | 0.058 | 0.256 | 0.430 | 0.001 |

**Burrowed material.**

| day | PDA + V (CHC/Met) | PDA + CHC/Met | PDA + V (Gal/Met) | PDA + Gal/Met | Tel + PDA |
|---|---|---|---|---|---|
| **0** | 0.027 | 0.000 | 0.054 | 0.085 | 0.022 |
| **1** | 1.000 | 0.311 | 1.000 | 1.000 | 1.000 |
| **2** | 1.000 | 0.305 | 1.000 | 1.000 | 1.000 |
| **4-8** | 0.197 | 0.003 | 0.377 | 0.032 | 1.000 |
| **18-19** | 1.000 | 0.538 | 0.758 | 0.025 | 1.000 |
| **34-35** | 1.000 | 0.080 | 0.405 | 1.000 | 1.000 |

**Nesting score.**

| day | PDA + V (CHC/Met) | PDA + CHC/Met | PDA + V (Gal/Met) | PDA + Gal/Met | Tel + PDA |
|---|---|---|---|---|---|
| **0** | 1.000 | 1.000 | 1.000 | 0.486 | 1.000 |
| **1** | 0.588 | 0.395 | 1.000 | 1.000 | 1.000 |
| **2** | 0.588 | 1.000 | 1.000 | 1.000 | 1.000 |
| **4-8** | 1.000 | 0.751 | 1.000 | 1.000 | 1.000 |
| **18-19** | 1.000 | 1.000 | 1.000 | 1.000 | 1.000 |
| **34-35** | 1.000 | 1.000 | 1.000 | 1.000 | 1.000 |

**Distress score.**

| day | PDA + V (CHC/Met) | PDA + CHC/Met | PDA + V (Gal/Met) | PDA + Gal/Met | Tel + PDA |
|---|---|---|---|---|---|
| **0** | - | 1.000 | 0.003 | 0.000 | 0.000 |
| **1** | - | 1.000 | 1.000 | 1.000 | 1.000 |
| **2** | - | 1.000 | 1.000 | 1.000 | 1.000 |
| **4-8** | - | 1.000 | 1.000 | 0.056 | 1.000 |
| **18-19** | - | 1.000 | 1.000 | 1.000 | 1.000 |
| **34-35** | - | 1.000 | 1.000 | 1.000 | 1.000 |

#### Table S2 – Pancreatitis

**Body weight change.**

| day | Panc + CR (miRNA-21 inh.) | Panc + miRNA-21 inh. |
|---|---|---|
| **2** | 0.012 | 0.009 |
| **16** | 0.000 | 0.003 |
| **30** | 0.001 | 0.003 |

**Burrowed material.**

| day | Panc + CR (miRNA-21 inh.) | Panc + miRNA-21 inh. |
|---|---|---|
| **2** | 0.000 | 0.028 |
| **16** | 0.016 | 0.152 |
| **30** | 0.003 | 0.152 |

**Nesting score**

| day | Panc + CR (miRNA-21 inh.) | Panc + miRNA-21 inh. |
|---|---|---|
| **2** | 0.155 | 0.091 |
| **16** | 0.040 | 0.000 |
| **30** | 0.040 | 0.003 |

**Distress score.**

| day | Panc + CR (miRNA-21 inh.) | Panc + miRNA-21 inh. |
|---|---|---|
| **2** | 1.000 | 1.000 |
| **16** | 0.351 | 0.472 |
| **30** | 0.126 | 1.000 |

#### Table S3 – $CCl_4$

**Body weight change.**

| day | $CCl_4$ + V (MCC950) | $CCl_4$ + MCC950 |
|---|---|---|
| **0** | 0.247 | 0.780 |
| **4** | 0.247 | 0.795 |
| **18** | 0.065 | 0.780 |
| **39** | 0.065 | 0.795 |

**Burrowed material.**

| day | $CCl_4$ + V (MCC950) | $CCl_4$ + MCC950 |
|---|---|---|
| **0** | 0.192 | 1.000 |
| **4** | 0.989 | 0.984 |
| **18** | 0.625 | 0.984 |
| **39** | 0.470 | 1.000 |

**Nesting score.**

| day | $CCl_4$ + V (MCC950) | $CCl_4$ + MCC950 |
|---|---|---|
| **0** | 0.002 | 0.004 |
| **4** | 0.469 | 0.644 |
| **18** | 0.000 | 0.004 |
| **39** | 0.005 | 0.004 |

**Distress score.**

| day | $CCl_4$ + V (MCC950) | $CCl_4$ + MCC950 |
|---|---|---|
| **0** | - | 1.000 |
| **4** | - | 1.000 |
| **18** | - | 0.455 |
| **39** | - | 1.000 |

#### Table S4 – Bile Duct Ligation

**Body weight change.**

| day | BDL + V (MCC950) | BDL MCC950 |
|---|---|---|
| **0** | 0.042 | 0.023 |
| **1** | 0.042 | 0.005 |
| **4** | 0.003 | 0.001 |
| **13** | 0.000 | 0.000 |

**Burrowed material.**

| day | BDL + V (MCC950) | BDL MCC950 |
|---|---|---|
| **0** | 0.002 | 0.005 |
| **1** | 0.085 | 0.007 |
| **4** | 0.044 | 0.035 |
| **13** | 0.000 | 0.001 |

**Nesting score.**

| day | BDL + V (MCC950) | BDL MCC950 |
|---|---|---|
| **0** | 0.001 | 0.000 |
| **1** | 0.061 | 0.005 |
| **4** | 0.151 | 0.046 |
| **13** | 0.110 | 0.005 |

**Distress score.**

| day | BDL + V (MCC950) | BDL MCC950 |
|---|---|---|
| **0** | 0.014 | 0.042 |
| **1** | 0.005 | 0.011 |
| **4** | 0.001 | 0.007 |
| **13** | 0.000 | 0.000 |
